# The Pharmacodynamic-Toxicodynamic Relationship of AUC and CMAX in Vancomycin Induced Kidney Injury in an Animal Model

**DOI:** 10.1101/2020.08.27.270793

**Authors:** Sean N. Avedissian, Gwendolyn Pais, Jiajun Liu, J. Nicholas O’Donnell, Thomas P. Lodise, Michael Neely, Walter C. Prozialeck, Peter C. Lamar, Leighton Becher, Marc H. Scheetz

## Abstract

**Background:** Vancomycin induces exposure-related acute kidney injury. However, the pharmacokinetic-toxicodynamic (PK-TD) relationship remains unclear.

**Methods:** Sprague-Dawley rats received IV vancomycin doses of 300mg/kg/day and 400mg/kg/day, divided once, twice, thrice or 4xdaily (i.e., QD, BID, TID or QID) over 24-hours. Up to 8-samples were drawn during the 24-hour dosing period. Twenty-four-hour urine was collected and assayed for kidney injury molecule-1 (KIM-1). Vancomycin was quantified via LC-MS/MS. Following terminal sampling, nephrectomy and histopathologic analyses were conducted. PK analyses were conducted using Pmetrics. PK exposures (i.e. AUC_0-24h_, CMAX_0-24h_,) were calculated for each rat, and PK-TD relationships were discerned.

**Results:** A total of 53-rats generated PK-TD data. A 2-compartment model fit the data well (Bayesian observed vs. predicted concentrations, R^2^=0.96). KIM-1 values were greater in QD and BID groups (P-values: QD vs TID:<0.002, QD vs QID:<0.004, BID vs TID:<0.002, and BID vs QID:<0.004). Exposure–response relationships were observed between KIM-1 vs CMAX_0–24h_ and AUC_0-24h_ (R^2^□=□ 0.7 and 0.68). Corrected Akaike’s information criterion showed CMAX_0-24h_ as most predictive PK-TD driver for vancomycin-induced kidney injury (VIKI) (−5.28 versus −1.95).

**Conclusions:** While PK-TD indices are often inter-correlated, maximal concentrations and fewer doses (for the same total daily amount) resulted in increased VIKI in our rat model.

## Introduction

Vancomycin was approved for clinical use in 1958 and is still one of the most commonly used antibiotics in the hospital setting because of its activity against methicillin-resistant *Staphylococcus aureus* (MRSA).[1] The initial pharmacokinetic/pharmacodynamic (PK/PD) efficacy studies performed in neutropenic mouse models demonstrated that the exposures calculated as area under the curve (AUC) divided by organism minimum inhibitory concentration (MIC) explained efficacy.[2, 3] Indeed, the 2020 vancomycin guidelines now recommend AUC monitoring to maximize efficacy for *S.aureus* infections.[4] The guidelines also noted that AUC monitoring and tighter control of vancomycin exposures may result in less kidney injury.

The AUC therapeutic window for vancomycin has been described in human and animal studies. A prospective clinical trial demonstrated that AKI increased above a 24 hour-AUC of 515 mg*h/L and that efficacy did not increase for MRSA bloodstream treatments above these exposures.[5] This ceiling threshold is consistent with other clinical data; a meta-analysis that included this study suggested a threshold AUC of 650 mg*h/L[6]. Preclinical rat studies have also found a consistent target, an AUC threshold of 482.2 has predicted 90% of maximal kidney injury biomarker response.[7] Thus, there is considerable evidence to support AUC as useful predictor of kidney injury, and thresholds are similar between humans and rats. It is still unknown if AUCs or maximal concentrations (Cmax) drives the toxicodynamic relationship for kidney injury. Herein, we present the results of dose fractionation experiments to better understand the PK/TD driver of vancomycin-induced kidney injury (VIKI).

## Materials and Methods

This PK/TD study was conducted at Midwestern University in Downers Grove, IL. All study methods were approved by the Institutional Animal Care and Use Committee (IACUC; Protocol #2295) and conducted in compliance with the National Institutes of Health Guide for the Care and Use of Laboratory Animals.[8]]

### Experimental design and animals

Experimental methods and design were similar to those described previously.[7] Male Sprague-Dawley rats (N=53, approximately 8-10 weeks old, mean purchase weight 310g) were housed individually in a light and temperature-controlled room for the duration of the study and allowed free access to water and food. Rats (n=5-9 per dosing protocol) were administered IV injections of clinical-grade vancomycin (n=48) in normal saline (NS) or NS only (n=5, control) as previously described.[7] In brief, rats were placed into a treatment or control group (treatment receiving vancomycin or normal saline, respectively). Vancomycin-treated rats received total daily doses of 300, or 400 mg/kg as either a thrice or four times daily divided dose over 24 hours (e.g., 300 mg/kg was given as a thrice injection [100 mg/kg three times daily] or as 75 mg/kg four times daily for a total of 24 hours). Previous animal data from our lab comprised the 300 and 400 mg/kg/day daily and twice daily cohorts (i.e., QD and BID) [7]. A complete animal dosing flow chart can be found in supplemental Figure 1. The 300-400 mg/kg/day dosing range was chosen based on the known nephrotoxic effect observed in our previous IV study [7] and to span the higher end of the clinical allometric range. For example, the clinical kidney injury threshold of ≥ 4 grams/day in a 70-kg patient (i.e., 57 mg/kg/day in humans) scales allometrically to 350 mg/kg in the rat. [9, 10] Data were analyzed for all animals that were included in the protocol.

### Blood and urine sampling

Surgical catheters were implanted 24 hours prior to protocol initiation. Blood samples were drawn from a single right-side internal jugular vein catheter, and dosing occurred via the left-side internal jugular vein catheter. A maximum of 8 samples per animal were obtained and scheduled at 0, 15, 30, 60, 120, 240, 750 and 1440 min post first dose for the once daily and twice daily dosing treatment protocol. The thrice dosing treatment protocol animals were sampled at 0, 15, 30, 60, 120, 240, 480, and 504 min. The QID daily dosing treatment protocol animals were sampled at 0, 15, 30, 60, 240, 360, 384 and 1094 min. Each sample (0.25 mL aliquot) was replaced with an equivalent volume of NS to maintain euvolemia. Blood samples from vancomycin-treated animals were immediately transferred to a disodium ethylenediaminetetraacetic acid (EDTA, [Sigma-Aldrich Chemical Company, Milwaukee WI]) treated microcentrifuge tube and centrifuged at 3000 rpm for 10 minutes. Plasma supernatant was collected and stored at −80°C for batch sample analysis.

Following the 2 hour blood sample, animals were placed in metabolic cages for urine collection (Nalgene, catalogue # 650-0350, Rochester, NY) for the remainder of the 24 hour study (with the exception that they were briefly removed for scheduled blood samples and vancomycin doses). Urine volume was measured at 12 and 24 hours. Urine was centrifuged at 400 x *g* for 5 minutes, and the supernatant was stored at −80°C until batch analysis.

### Chemicals and reagents

Animals were administered clinical grade vancomycin hydrochloride for injection (Lot#: 591655DD) obtained commercially (Hospira, Lake Forrest, IL). All solvents were of liquid chromatography-tandem mass spectrometry (LC-MS/MS) grade. For LC-MS/MS assay purposes, vancomycin hydrochloride, United States Pharmacopeia was used (Enzo Life Science, Farmingdale, NY) with a purity of 99.3%. Polymyxin B (Sigma-Aldrich, St. Louis, MO), acetonitrile, and methanol were purchased from VWR International (Radnor, PA). Formic acid was obtained from Fischer Scientific (Waltham, MA). Frozen, non-medicated, non-immunized, pooled Sprague-Dawley rat plasma (anticoagulated with disodium EDTA) was used for calibration of standard curves (BioreclamationIVT, Westbury, NY).

### Determination of vancomycin concentrations in plasma

Plasma concentrations of vancomycin were quantified with LC and column conditions similar to those used in our previous report [7]. The lower limit of quantification was 0.25 mg/L. Precision was <8.6% for all measurements, including intra- and inter-day assay measurements. Greater than 92% of the analyte was recovered in all samples tested with an overall mean assay accuracy of 100%. Any samples measuring above the upper limit of quantification were diluted per standard protocol and requantified.

### Determination of urinary biomarkers of AKI

Urine samples were analyzed in batch to determine concentrations of KIM-1. Microsphere-based Luminex X-MAP technology was used for the determination of all biomarker concentrations, as previously described. [11, 12] Urine samples were aliquoted into 96-well plates supplied with MILLIPLEX^®^ MAP Rat Kidney Toxicity Magnetic Bead Panels 1 and 2 (EMD Millipore Corporation, Charles, MO), prepared and analyzed according to the manufacturer’s recommendations.

### Histopathology Kidney Scoring

Following terminal blood sampling, bilateral nephrectomy was performed under anesthesia as previously described [13]. Briefly, kidneys were washed in cold isotonic saline and preserved in 10% formalin solution for histologic examination. Histopathologic analyses were conducted by IDEXX BioAnalytics (Westbrook, Maine). Pathologists only received access to nominal dosing group assignment. Scoring was conducted according to the Critical Path Institute’s Predictive Safety Testing Consortium Nephrotoxicity Working Group’s histologic injury lexicon which utilizes a 0-5 point ordinal scale (25). This scoring system assigns higher scores to increasing levels of damage (0, no evidence of damage; 1, minimal; 2, mild; 3, moderate; 4, marked; 5, severe/massive) and has been validated previously (17, 25). The composite score for each animal was calculated as the highest ordinal score for histopathologic changes at any kidney site (25).

### Vancomycin pharmacokinetic model and exposure determination

We employed the Bayesian priors from our previously published pharmacokinetic model [7] to generated Bayesian posteriors for all N= 53 animals reported in this manuscript. Pharmacokinetic analyses were completed using the Pmetrics package version 1.5.0 (Los Angeles, CA) for R version 3.2.1 (R Foundation for Statistical Computing, Vienna, Austria)[14, 15] with model assessment as previously described [7].

### Estimation of PK exposure profiles and statistical analysis

The pharmacokinetic model was utilized to obtain median maximum a posteriori probability (MAP) Bayesian vancomycin plasma concentration estimates at 12-minute intervals over the 24 hour study period, generated from each animal’s measured vancomycin concentrations, exact dose, and dosing schedule. Bayesian posteriors for each animal were used to determine exposures over the 24-hour time period (i.e., AUC_0-24h_, CMAX_0-24h_). The pharmacokinetic value CMAX_0-24h_ were calculated using ‘makeNCA’ within Pmetrics (Los Angeles, CA, USA) [14, 16]. The highest Bayesian posterior concentration was determined to be each individual animal’s CMAX_0-24h_. Twenty-four hour exposure, as measured by AUC_0-24h_, was calculated using the trapezoidal rule within the Pmetrics command ‘makeAUC’ [14, 16]. Cumulative CMAX_0-24h_ was also calculated for the dose fractionation groups (i.e., BID CMAX multiplied by 2, TID CMAX multiplied by 3, QID CMAX multiplied by 4) and standardized to mg to allow comparison and assess successfulness of varying CMAX_0-24h_ while holding AUC_0-24h_ constant.

### Association of PK measures with urinary AKI biomarker KIM-1

Pharmacokinetic exposure estimates were assessed for relationships with KIM-1 using GraphPad Prism version 7.02 (GraphPad Software Inc., La Jolla, CA). PK/TD exposure-response relationships with KIM-1 were evaluated using Spearman’s rank correlation coefficient. Hill-type functions and log transformations of variables were employed to explore the relationship between PK exposures and KIM-1. Correlation coefficients (R^2^) and corrected Akaike information criterions (AICc) were calculated and compared between exposures metrics (CMAX vs. AUC) to evaluate overall fit. KIM-1 was the primary biomarker of interest given specificity for proximal tubule damage and identification of VIKI in the rat model [7, 17, 18].

### Statistical analysis for between treatment group comparisons

Statistical analysis for between treatment group comparisons was performed using Intercooled Stata, version 14.2 (College Station, TX: StataCorp LP.). PK exposure measurements, i.e. AUC_0-24h_, CMAX_0-24h_, were compared across vancomycin total daily dose and dosing frequency groups for 300 mg/kg/day and 400 mg/kg/day. Log transformations were employed as needed. Differences between treatment groups for KIM-1 and standardized CMAX were visualized with LOWESS regression and compared with the Kruskal-Wallis Dunn pairwise comparison with the Bonferroni adjustment. All tests were two-tailed, with an *a priori* level of statistical significance set at an alpha of 0.05.

## Results

### Characteristics of animal cohort

All 48 animals from the 300 mg/400 mg total daily dose cohorts contributed pharmacokinetic model data (5 controls were not included in model since, by design, they did not have quantifiable vancomycin levels). Mean baseline weights were not significantly different between controls and vancomycin dosing protocol animals (307.4 g versus 313.6 g, P=0.49). Overall 24-hour urine output was significantly different between controls and the vancomycin treated animals (4.2 versus 16.11 mL, P<0001). Lastly, median histopathology scores differed numerically though not statistically significantly between controls and the entire vancomycin treated cohort (1 versus 2, P=0.088). However, median KIM-1 values were significantly different between controls and vancomycin dosing protocol animals (0.63 ng/mL versus 6.169 ng/mL, p<0.001).

In stratified dosing group analyses (i.e. 300 mg/kg/day and 400 mg/kg/day), there was a significant difference in median KIM-1 between fractionation schemes. In the 300 mg/kg/day group, median KIM-1 values were significantly different in the daily vs TID (10.7 ng/mL vs. 2.3 ng/mL, P=0.03) and daily vs QID (10.70 ng/mL vs. 1.46 ng/mL, P<0.01). In the 400 mg/kg/day group, median KIM-1 values only differed in the BID vs QID (13.3 ng/mL vs. 4 ng/mL) dose fractionation group (P-value: <0.001) A complete pairwise comparisons can be found in Table 1.

**Table 1:** Biomarker KIM-1 summary for vancomycin treated animals by dose fractionation

| **300 mg/kg/day animals** | **Daily**<br>**(N=5)** | **BID**<br>**(N=5)** | **TID**<br>**(N=6)** | **QID**<br>**(N=6)** | **P-value** |
| --- | --- | --- | --- | --- | --- |
| Median KIM-1<br>(ng/mL)<br>(IQR) | 10.70* **<br>(9.8-12.3) | 9.5<br>(7.7-10.3) | 2.3*<br>(1.43-4.9) | 1.46**<br>(1.2-2.5) | *0.03<br>**<0.01 |
| **400 mg/kg/day animals** | **Daily**<br>**(N=5)** | **BID**<br>**(N=6)** | **TID**<br>**(N=9)** | **QID**<br>**(N=6)** | **P-value** |
| Median KIM-1<br>(ng/mL)<br>(IQR) | 10.5<br>(6.6-14.4) | 13.3*<br>(13-16.5) | 4*<br>(2.9-4.6) | 6.4<br>(4.8-8.3) | *<0.001 |

Abbreviations: KIM-1= kidney injury molecule-1, IQR= interquartile range, BID= twice daily, TID= three times daily, QID=four times daily

### Vancomycin PK models, parameter estimates and exposures

Median (CV%) parameter values for the pharmacokinetic model for Ke, V, KCP and KPC were: 0.7 hr^−1^ (60.81), 0.07 L (76.69), 1.42 hr^−1^ (136.87), 1.52 hr^−1^ (160.86), respectively. Model predictive performance demonstrated observed versus Bayesian predicted concentrations, bias, imprecision (i.e., bias-adjusted mean weighted squared prediction error) and the coefficient of determination (R^2^) of 0.101 mg/L, 2.61 (mg/L)^2^, and 0.958 respectively (Figure 1). A complete concentration vs. time profile plot by dose fractionation for all animals in the 300 mg/kg/day and 400 mg/kg/day can be found in Figure 2. There were no significant differences found in CMAX standardized by fractionation and mg in all animals.

**Figure 1.**
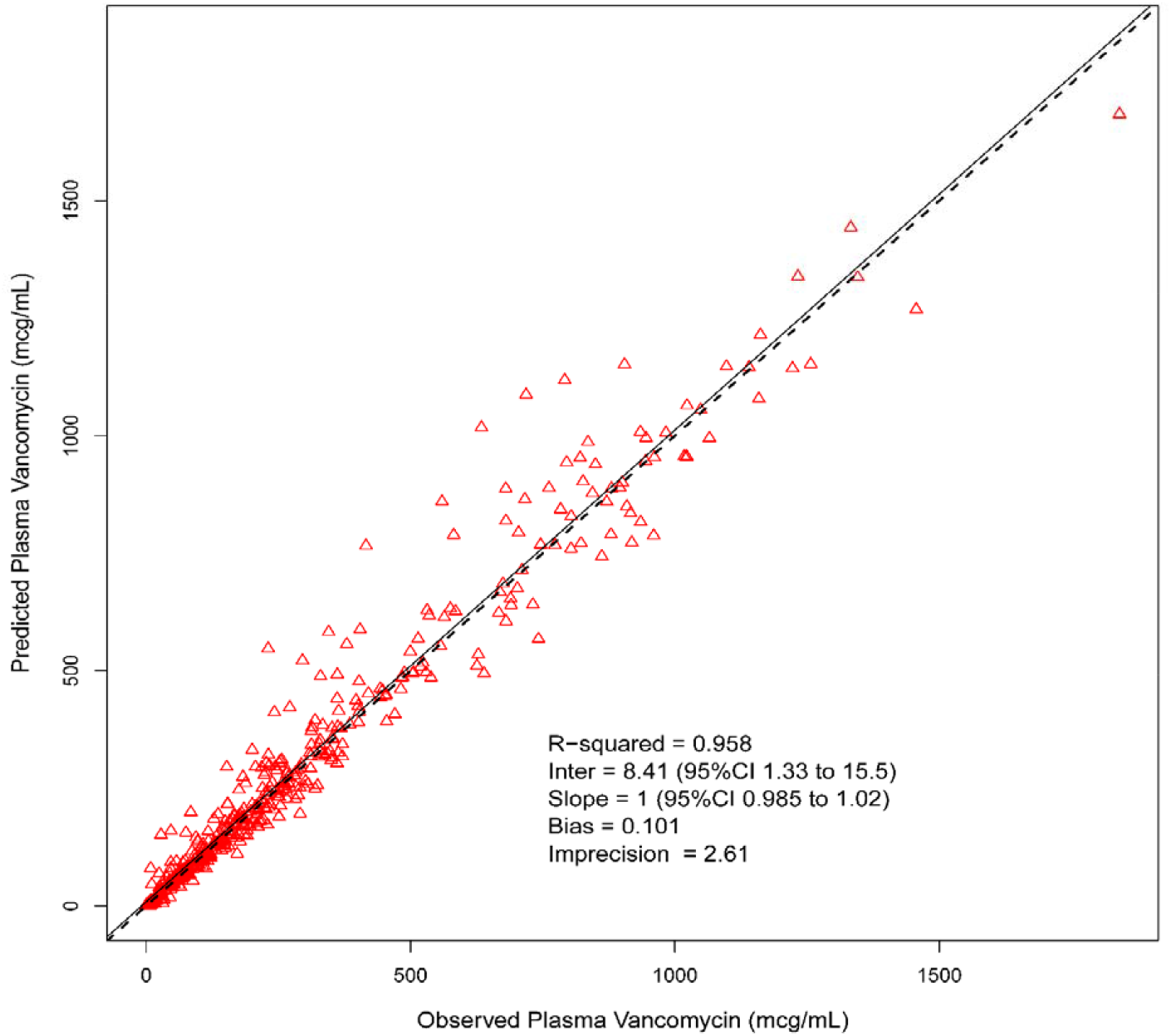
Best fit plot for Bayesian observed versus predicted plasma vancomycin concentrations utilizing the final 2-compartmental model

**Figure 2.**
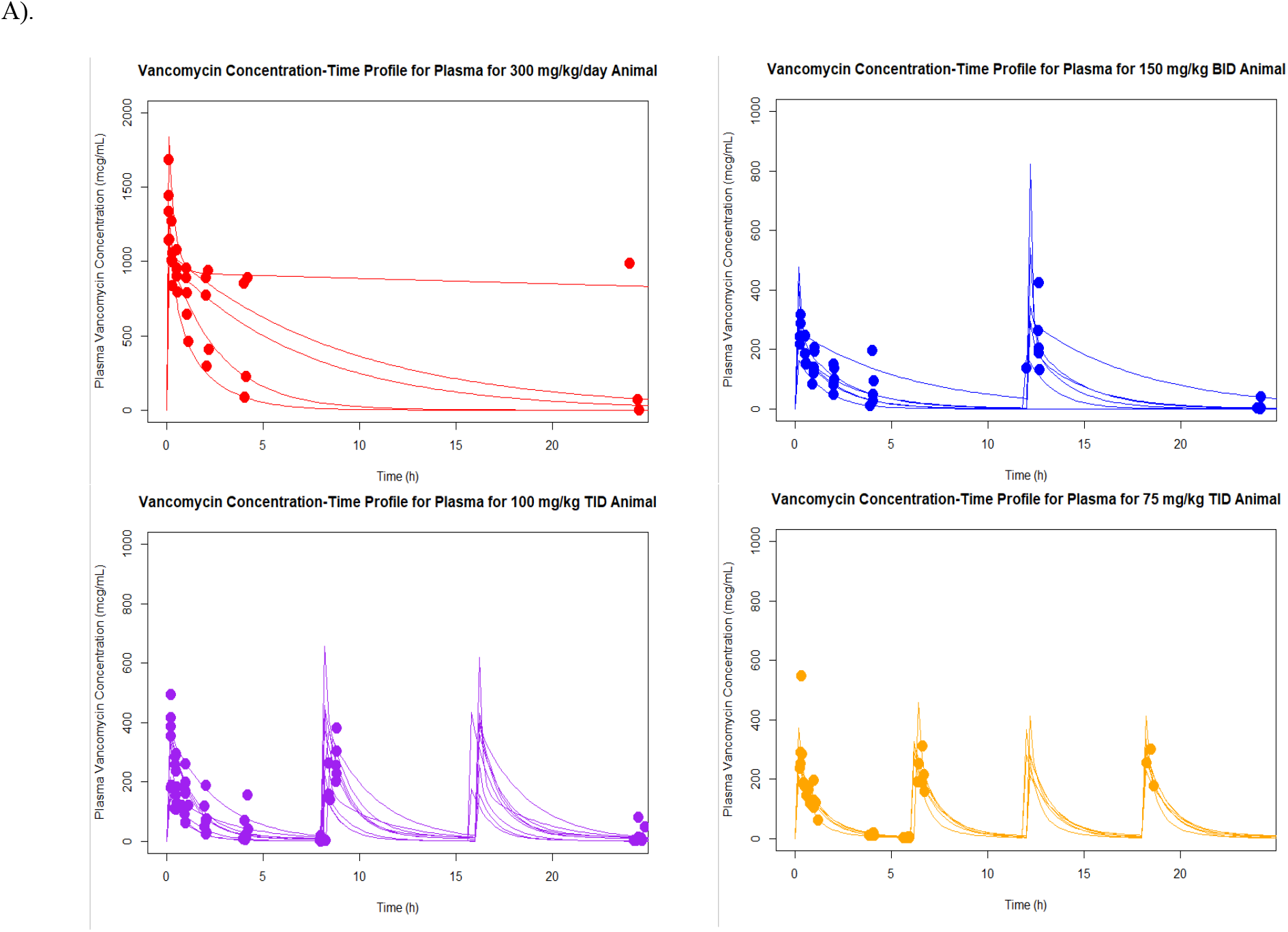
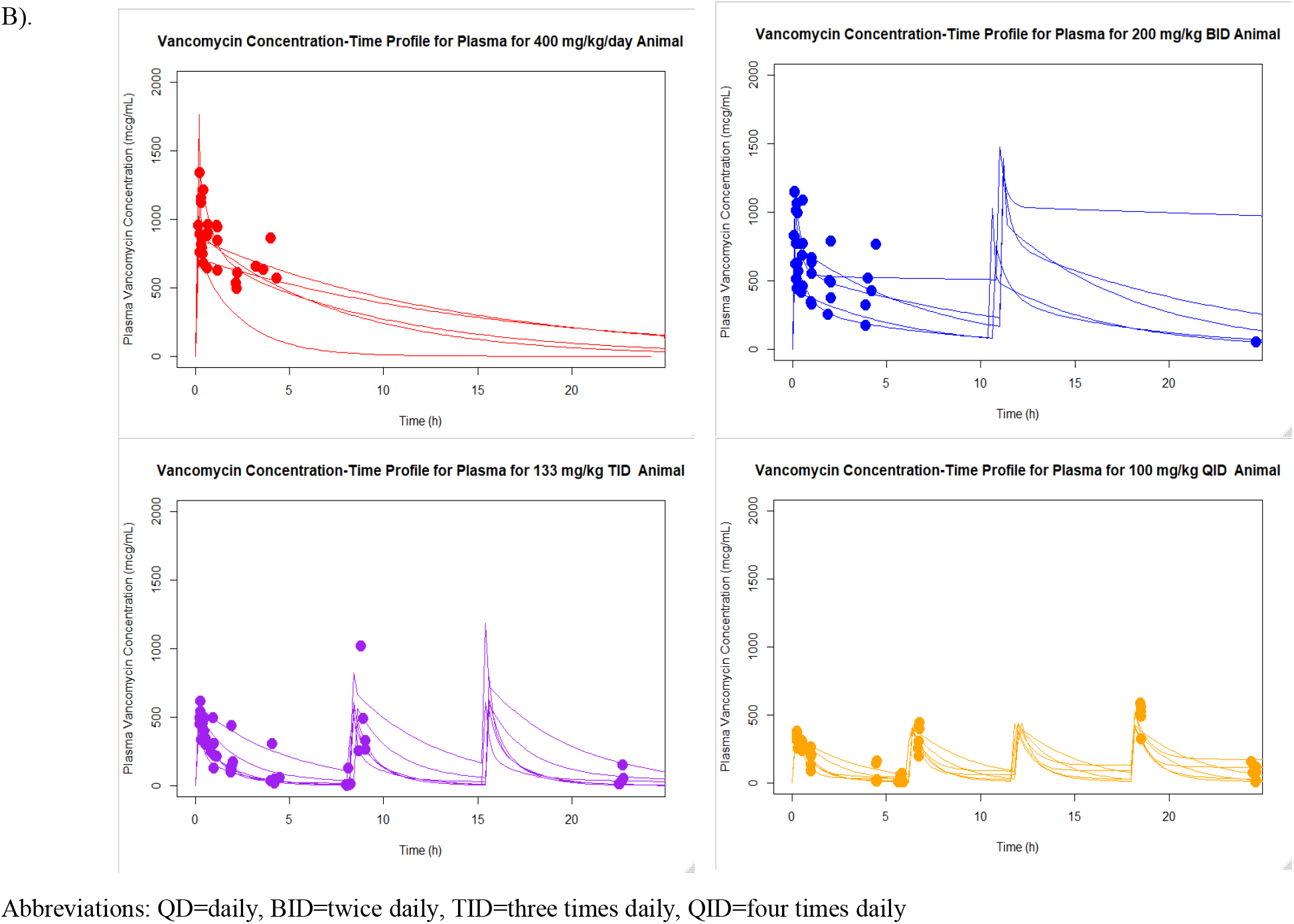
Concentration versus time plots for each dose fractionation group all animals (A) 300 mg/kg/day and (B) 400 mg/kg/day A).

### Exposure-Response Relationships

Four parameter Hill models best described the exposure:biomarker relationships. Exposure:biomarker relationships were found between AUC_0-24_ versus KIM-1 (R^2^=0.68) and CMAX_0-24_ versus KIM-1 (R^2^=0.7). Overall, AUC_0-24_ versus KIM-1 was slightly less predictive than CMAX_0-24_ versus KIM-1 as the CMAX_0-24_ model performed better based on AICc comparison (AICc=-5.28 vs. AICc= −1.95). All exposure-biomarker relationships are shown in Figure 3.

**Figure 3.**
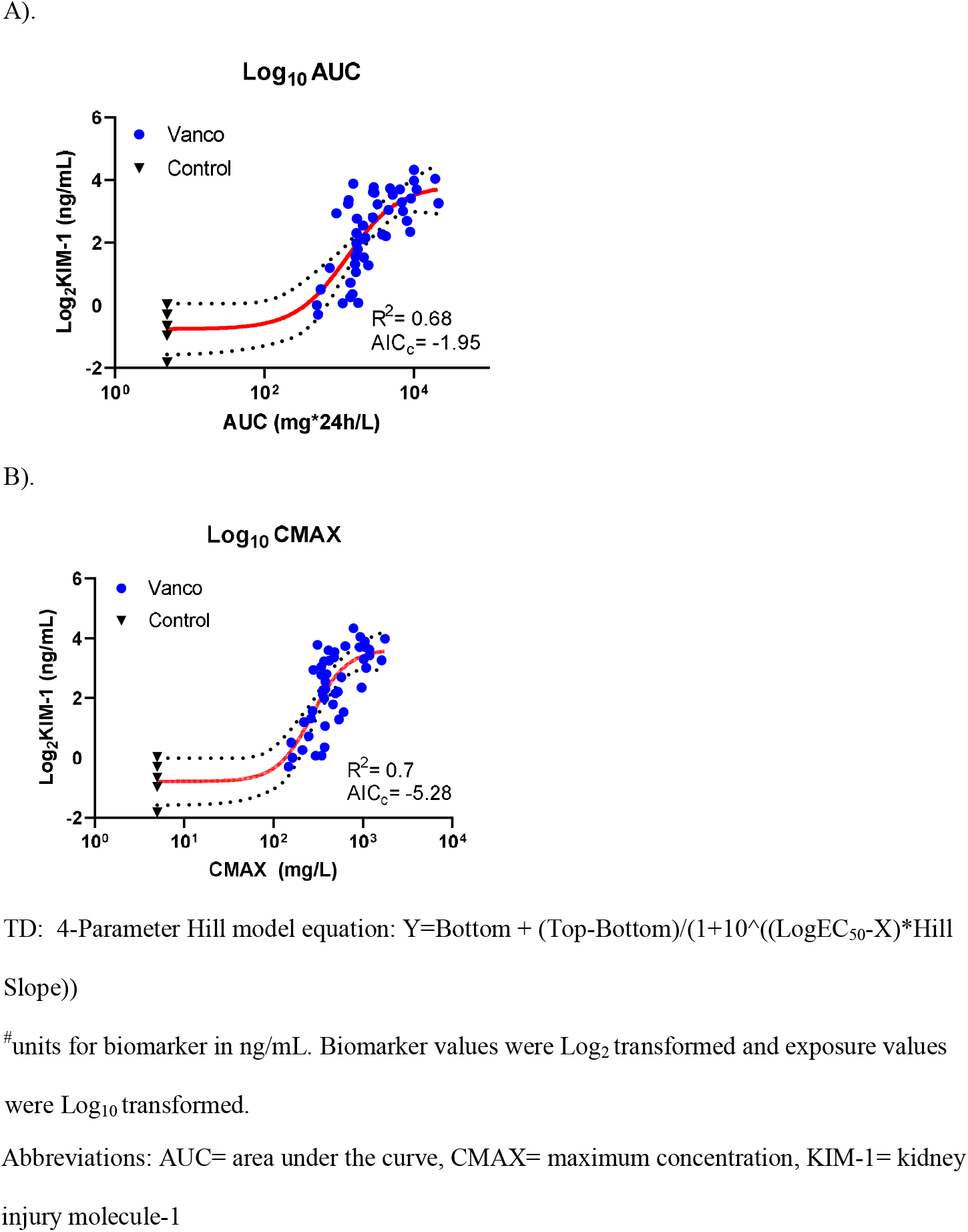
(A) AUC_0-24_ (mg*h/L), and (B) CMAX_0-24_ (mg/L) versus urinary biomarkers KIM-1^#^ (A, B, C) and OPN* (D, E, F) relationship by 4-Parameter Hill model fit

### Vancomycin Dose Fractionation vs Biomarker Relationships

The plot visually displays decreasing urinary KIM-1 concentrations as doses were increasingly fractionated between once and four times daily for both daily dose groups (i.e. 300 and 400 mg/kg/day). Pairwise comparison showed that there was a significant difference in KIM-1 between QD vs TID and QID groups and BID to TID and QID (P-values all <0.01). A visual representation of these trends and differences are shown in Figure 4.

**Figure 4.**
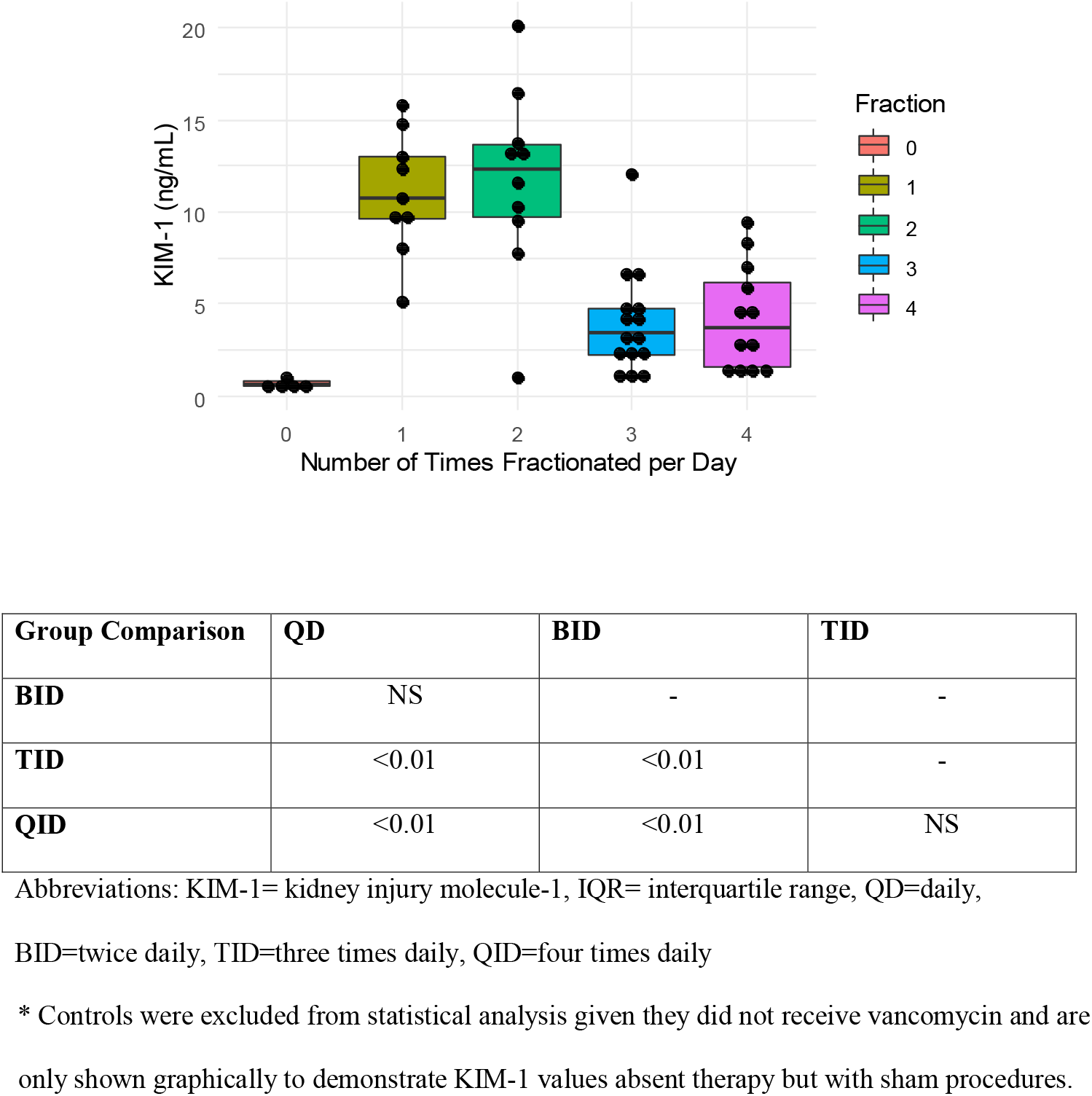
Dose Fractionation vs. KIM-1 Relationship for all Animals

## Discussion

This pre-clinical study provides continued evidence that VIKI is caused by high maximal vancomycin concentrations and elevated exposures. Importantly in our model, CMAX was marginally better than AUC in explaining toxicodynamic relationships. These findings, along with studies that have compared prolonged infusion vancomycin to standard intermittently infused vancomycin, suggest that giving the same total daily dose in fractionated fashion or with continuous infusion may improve kidney outcomes.[19–23] In our trial where we were able to fractionate doses, we identified that CMAX exhibited a slightly improved relationship with KIM-1 compared to AUC 4-parameter Hill model fit (CMAX verus KIM-1: R^2^ = 0.7, AIC_c_ = −5.28; AUC versus KIM-1: R^2^ =0.68, AIC_c_ = −1.95). Further, the fractionation scheme resulted in lower KIM-1. That is, when the daily dose was split into several doses, kidney injury was less. For the fractionation schemes, we demonstrated that CMAX remained constant between the groups (p=0.34) and that urinary KIM-1 was lower in TID and QID groups than QD and BID (P-values all <0.01).

The rat is a highly relevant pre-clinical model for drug induced acute kidney injury, as quantiative biomarkers that describe degree of injury are shared between humans and rats.[24, 25] Adequate blood sample volumes are possible in the rat, thus allowing for a richly sampled PK design and careful characterization of vancomyicn exposures in order to assess pharmacokinetic/toxicodynamic (PK/TD) relationships at the indivdual animal level. Further, PK/TD relationships are similar between this rat model and human VIKI. In both species, toxicity thresholds are ~500-600 mg/L*24hours.[5–7] KIM-1 was utilized as the surrogate of VIKI in this study as it is causally linked to histopathologic damage in V treated rats. KIM-1 is a specific marker of histopathologic proximal tubule injury in VIKI [7, 13]. KIM-1 has predicted histopathologic rise (P<0.001) and was the best predictor of histopathologic damage score ≥2 on each study day (i.e. day 1, 2, 3, and 6) as determined by ROC area. Quantitatively, every 1 ng/mL increase of KIM-1 increased the likelihood of a histopathologic score ≥ 2 by 1.3 fold (P<0.001).[5]

Our findings by fractionation group are similar to those from Konishi et al.’s study in Wistar rats. They found that creatinine clearance and superoxide dismutase were decreased in rats treated once daily vs. twice daily.[26] Our study is different in that we administered vancomycin intravenously and were additionally able to precisely estimate exposures for each rat in order to define the PK/TD relationships. Konishi et al. studied animals over 7 days while we studied a single day (as this is the PK/TD model that has best linked rat outcomes to human outcomes).[5, 7, 26] Their findings that vancomycin saturated similarly in the kidney between the two groups may suggest that outcomes will eventually converge toward equivocal if treatment is prolonged. That is, fractionating doses may provide benefit early; however, it will remain prudent to discontinue nephrotoxic drugs when they are not needed as human trials and animal models consistently show the kidney toxicity of vancomycin.[27]

Randomized studies will be necessary to discern if fractionating the daily dose of vancomycin ultimately improves outcomes for humans. Indeed, small clinical studies and meta-analyses have demonstrated that continuous infusions of vancomycin might result in less kidney injury than traditional intermittent infusions.[19, 21, 22] In theory, limiting the CMAX even for equivalent AUC exposures could improve the renal safety of vancomycin. The single prospective human trial that randomized patients to continuous infusion vancomycin (n=61) or traditional intermittent infusions (n=58) did not find a difference in renal outcomes between the groups. However the study was relatively underpowered to assess this outcome in the setting of a heterogenous study population receiving varied concomitant therapies.[28] A prolonged vancomycin administration has also been considered in recent national guidelines which concluded with moderate evidence that the risk of developing nephrotoxicity with continuous infusion appears to be similar or lower than that with intermittent dosing.[4]

There are several limitations to this study. First, this study was limited to 24 hour dosing for our dose fractionation protocol. However as previously noted, elevations in biomarkers have already been linked to histopathologic damage within this time period [29]. Second, this employed allometric scaled doses that are known to result in toxicity, i.e. CMAX was not humanized. Additional studies will be needed to understand if the TD relationship found in this study is reproducible if CMAX is scaled to humanized values. It is notable that it is not possible to utilize standard practices of administering nephrotoxic agents to animals match human clearance when the outcome being assessed is AKI. Thus, continuous infusion may be the best way to parameter scale CMAX, and it is not clear if those studies will be more translational than the current approaches.

In summary, these data demonstrate that VIKI may be driven by CMAX_0-24_. These findings have clinical implications as dosing strategies may be able to dose fractionate a total daily vancomycin dose in efforts to maintain efficacy by maintaining AUC while decreasing toxicity. Further studies employing continuous infusion dosing strategies are warranted to further assess if administration scheme can mitigate toxicity.

## Acknowledgments

We kindly acknowledge the Core Facility at Midwestern University for access to the LCMS-MS.

---

**Supplemental Figure 1.**
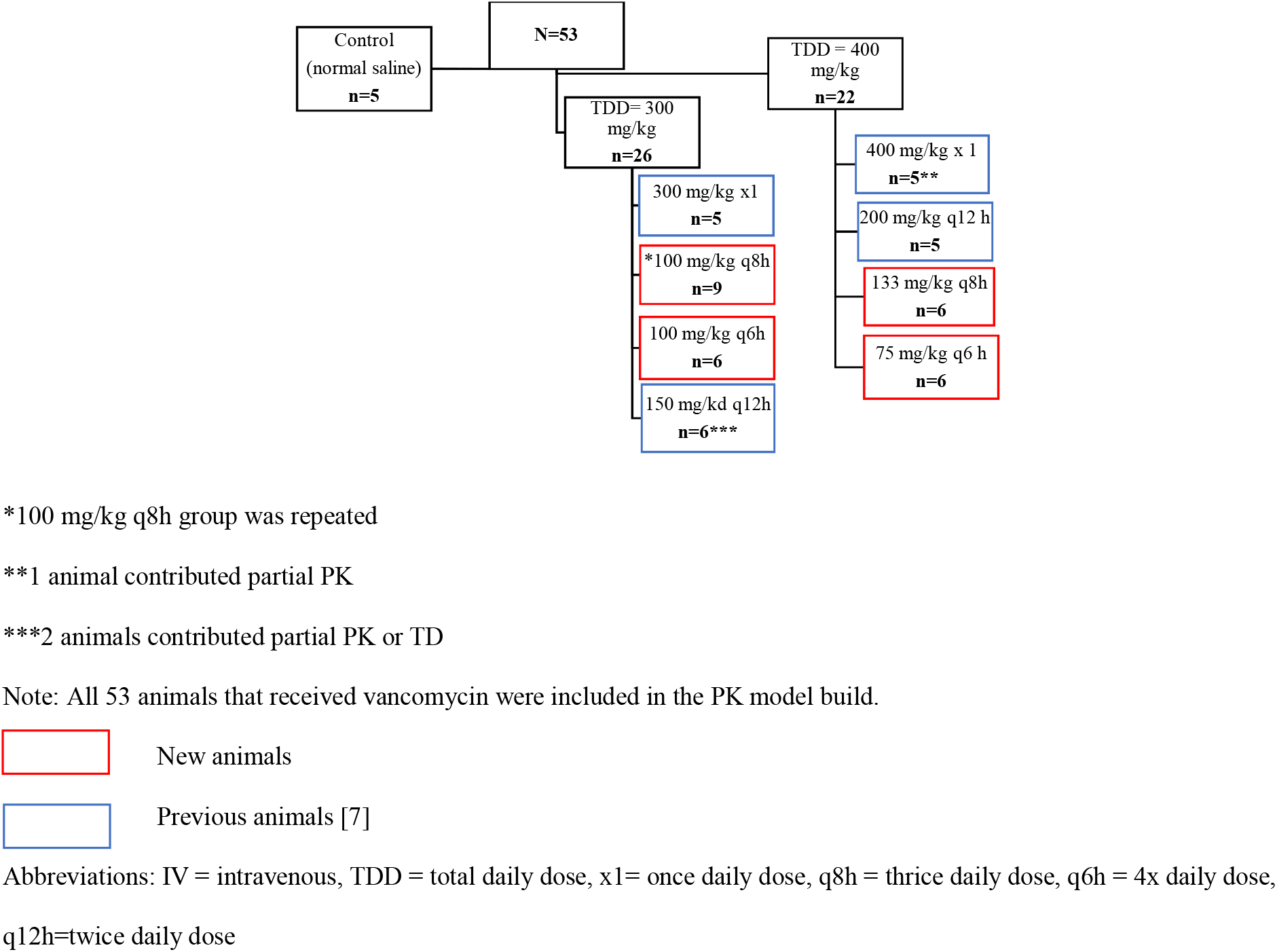
Animal dosing flow chart

